# Bisphosphonate drugs have actions outside the skeleton and inhibit the mevalonate pathway in alveolar macrophages

**DOI:** 10.1101/2021.08.09.455652

**Authors:** Marcia A. Munoz, Emma K. Fletcher, Oliver P. Skinner, Julie Jurczyluk, Esther Kristianto, Mark P. Hodson, Shuting Sun, Frank H. Ebetino, David R. Croucher, Philip M. Hansbro, Jacqueline R. Center, Michael J. Rogers

**Author notes:** Address for correspondence: Prof Mike Rogers, Garvan Institute of Medical Research, 384 Victoria St, Darlinghurst, Sydney, NSW 2010, Australia.

## Abstract

Bisphosphonates drugs target the skeleton and are used globally for the treatment of common bone disorders. Nitrogen-containing bisphosphonates act by inhibiting the mevalonate pathway in bone-resorbing osteoclasts but, surprisingly, also appear to reduce the risk of death from pneumonia. We overturn the long-held belief that these drugs act only in the skeleton and show that a fluorescently-labelled bisphosphonate is internalised by alveolar macrophages and peritoneal macrophages *in vivo*. Furthermore, a single dose of a nitrogen-containing bisphosphonate (zoledronic acid) in mice was sufficient to inhibit the mevalonate pathway in tissue-resident macrophages, causing the build-up of a mevalonate metabolite and preventing protein prenylation. Importantly, one dose of bisphosphonate enhanced the immune response to bacterial endotoxin in the lung and increased the level of cytokines and chemokines in bronchoalveolar fluid. These studies suggest that bisphosphonates, as well as preventing bone loss, may boost immune responses to infection in the lung and provide a mechanistic basis to fully examine the potential of bisphosphonates to help combat respiratory infections that cause pneumonia.

## INTRODUCTION

Nitrogen-containing bisphosphonates (N-BPs) are a class of bone-seeking drugs used worldwide as the front-line treatment for disorders of excessive bone resorption such as post-menopausal osteoporosis and cancer-associated bone disease (Russell, 2011). By virtue of their avidity for calcium ions, N-BPs bind rapidly to the skeleton, where they are internalised by bone-degrading osteoclasts (Russell et al., 2008, Rogers et al., 2020). Intracellularly, N-BPs disable osteoclast function by inhibiting the enzyme farnesyl diphosphate (FPP) synthase in the mevalonate pathway (van Beek et al., 1999, Dunford et al., 2001), thereby preventing the post-translational prenylation of small GTPase proteins necessary for osteoclast function (Luckman et al., 1998, Fisher et al., 1999).

There is increasing evidence that N-BP drugs, such as zoledronic acid (ZOL), have benefits beyond preventing bone loss (Center et al., 2020) and, unexpectedly, N-BP therapy has recently been linked to reduced risk of mortality from pneumonia (Colon-Emeric et al., 2010, Sing et al., 2020). In a randomised, controlled trial of >2,000 hip fracture patients, ZOL therapy reduced the risk of death by 28% compared to placebo infusion (Lyles et al., 2007). Retrospective analysis also suggested that ZOL-treated patients were less likely to die from pneumonia than placebo-treated subjects (Colon-Emeric et al., 2010). Recently, a “real-world” population-based, observational study of hip fracture patients aged 50 years or above also showed a significant reduction in risk of pneumonia and pneumonia mortality in hip fracture patients that had received N-BP therapy, compared to no treatment or other osteoporosis medications (Sing et al., 2020). Similar findings were reported in the post hoc analysis of a randomised controlled trial of ZOL in women over the age of 65 years (Reid et al., 2021). Pneumonia is the most frequent cause of admission to ICU and a study of long term patients in respiratory ICU revealed a significant reduction in mortality in people treated with the N-BP pamidronate compared to those without treatment (Schulman et al., 2016). Furthermore, in a retrospective cohort study of ICU subjects, we showed a 59% reduction in mortality in patients treated with bisphosphonate prior to hospitalisation (Lee et al., 2016). However, the mechanisms underlying the surprising beneficial effects of these drugs on pneumonia and ICU patients are unknown.

Globally, respiratory diseases constitute the most common cause of death and thus, therapies that boost the immune response to common lung infections are urgently needed. Bacterial infections such as *Streptococcus pneumoniae*, *Haemophilus influenza, Chlamydia pneumoniae* and *Staphylococcus aureus* are the main cause of community-acquired pneumonia. Importantly, these pathogens also underlie severe complications of viral respiratory disease that can significantly increase morbidity and mortality. For example, influenza-related mortality is often associated with pneumonia caused by co- or secondary bacterial infection (Morris et al., 2017). In this study we debunk the long-held view that N-BP drugs act only in the skeleton and show that even a single dose of N-BP in mice is sufficient to affect tissue-resident macrophages, including lung alveolar macrophages (AMϕ), boosting their response to bacterial endotoxin.

## RESULTS AND DISCUSSION

### Systemically administered N-BP is internalised by tissue-resident macrophages outside the skeleton

We previously reported that cultured macrophages and tumour-associated macrophages *in vivo,* like osteoclasts, have the ability to internalise N-BP by endocytosis (Thompson et al., 2006, Junankar et al., 2015). In this study we used a fluorescently-labelled analogue of ZOL (AF647-ZOL) to determine whether tissue-resident macrophages are capable of internalising systemically administered N-BP in mice. Given the important role of AMϕ in lung homeostasis and the initial immune response to respiratory infection (Byrne et al., 2015, Crane et al., 2018), we focused on whether these cells are targeted by N-BP. To answer this question, animals were injected with a single intravenous (*i.v*.) dose of AF647-ZOL and cells collected by bronchoalveolar lavage (BAL) were analysed by flow cytometry. In the absence of immune challenge in mice, approximately 90% of cells in BAL were AMϕ (Suppl Fig 1a). Importantly, >98% of AMϕ (TCRβ^−^B220^−^CD11b^lo/−^ CD11c^hi^F4/80^+^) in BAL samples were clearly labelled with AF647-ZOL 4 hours after *i.v*. administration (Fig 1a–c).

**Figure 1.**
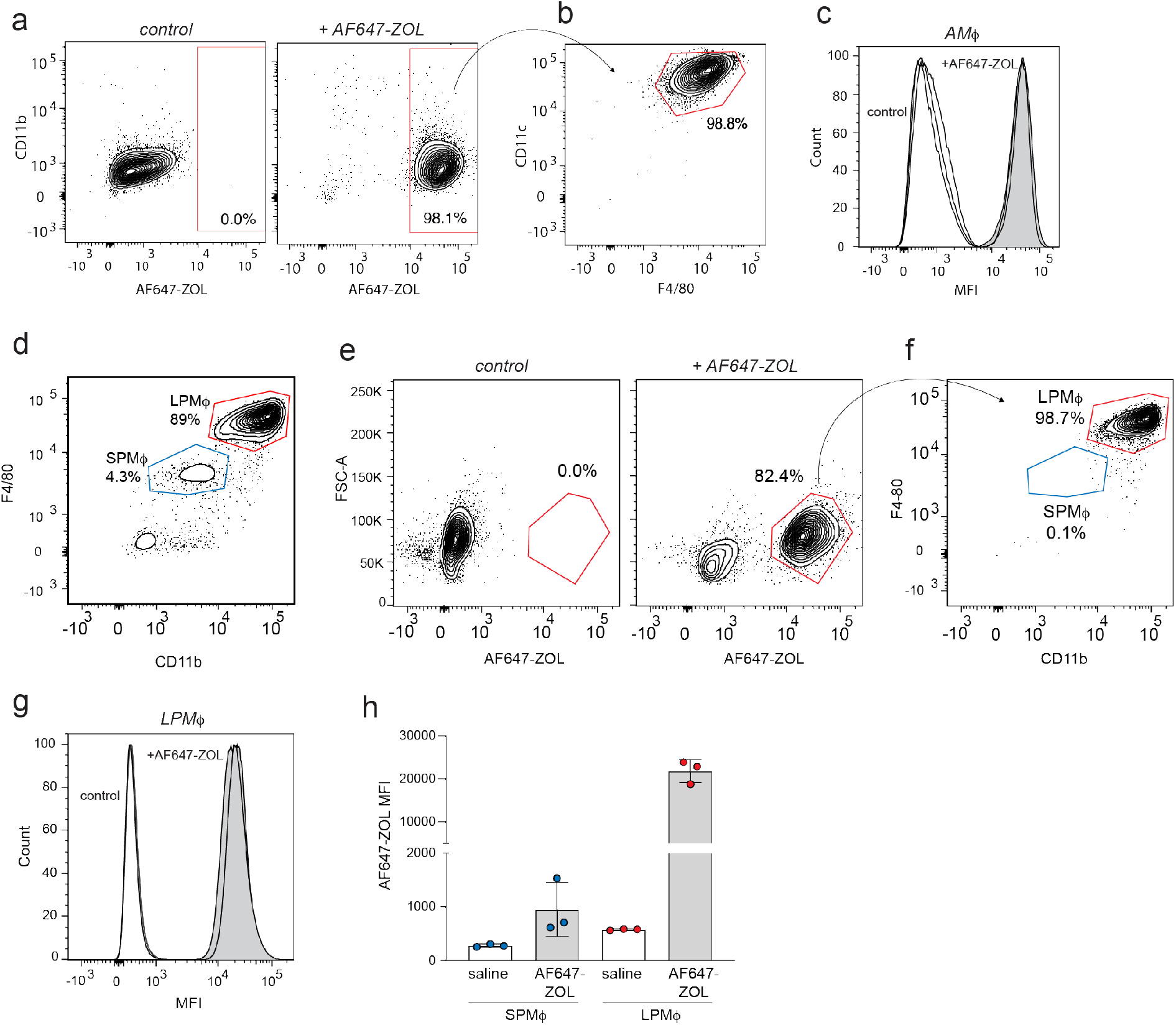
Bisphosphonate is internalised by tissue-resident macrophages *in vivo.* **a)** FACS plots showing the percentage of labelled single cells in BAL samples after one *i.v.* dose of AF647-ZOL, compared to saline-treated mice. **b)** AF647-ZOL positive cells were predominantly alveolar macrophages (AMϕ) i.e. B220^−^TCRb^−^, CD11c^+^F4/80^+^ singlets. **c)** Histograms show AF647-ZOL mean fluorescence intensity (MFI) of AMϕ in BAL samples from n=3 control mice (white) and n=3 AF647-ZOL-treated mice (grey). **d)** FACS plot illustrating the percentages of small peritoneal macrophages/SPMϕ (B220^−^TCRb^−^ singlets, CD11b^int^F4/80^int^) and large peritoneal macrophages/LPMϕ (B220^−^TCRb^−^ singlets, CD11b^hi^F4/80^hi^) in peritoneal lavage. **e)** Percentage of labelled peritoneal cells 4 hours after one *i.v.* injection of saline (left) or AF647-ZOL (right). **f)** The labelled cell population (AF647-ZOL^+^, 82.4%) in **e)** consists predominantly of CD11b^hi^F4/80^hi^ LPMϕ. **g)** Histograms show the mean fluorescence intensity (MFI) of LPMϕ from saline-(white) and AF647-ZOL-treated (grey) mice. **h)** Mean fluorescence intensity (AF647-ZOL MFI) values from SPMϕ and LPMϕ isolated from saline- or AF647-ZOL treated animals. Bars represent mean ± SD (n=3 mice per group in **g,h;** each symbol represents the measurement from an individual mouse). FACS plots in **a,b,d-f** are representative of 3 mice per group.

We also examined N-BP uptake in peritoneal macrophages (PMϕ). Under baseline conditions, 80% of cells obtained by peritoneal lavage (PL) consisted of PMϕ (B220^−^ TCRb^−^ Siglec-F^−^ Ly6G^−^ CD11b^+^ F4/80^+^), most of which were CD11b^hi^F4/80^hi^ large PMϕ, with a less abundant population of CD11b^+^F4/80^int^ small PMϕ (Ghosn et al., 2010) (Fig 1d, Suppl Fig 1a). Similar to BAL cells, approximately 80% of peritoneal cells incorporated AF647-ZOL after a single *i.v.* dose (Fig 1e), the majority of which (99%) were CD11b^hi^F4/80^hi^ large PMϕ (Fig 1f,g). In contrast, the CD11b^+^F4/80^int^ small PMϕ incorporated negligible amounts of fluorescently-labelled ZOL (Fig 1h). In addition to the well-described uptake of N-BP by osteoclasts in bone (Rogers et al., 2020, Coxon et al., 2008), these findings clearly demonstrate that N-BP can also be efficiently internalised *in vivo* by tissue-resident macrophages outside the skeleton, including AMϕ in the lung and LPMϕ in the peritoneal cavity.

### A single *i.v*. dose of N-BP is sufficient to inhibit the mevalonate pathway in alveolar and peritoneal macrophages

To examine whether tissue-resident macrophages can incorporate sufficient N-BP *in vivo* to have a pharmacologic effect, we analysed two biochemical outcomes that, together, are reliable features of intracellular N-BP action in cells (Rogers et al., 2020): (i) the cytoplasmic build-up of the upstream metabolite isopentenyl diphosphate (IPP) and its isomer dimethylallyl diphosphate (DMAPP) (Raikkonen et al., 2009); and ii) reduced production of the isoprenoid lipid geranylgeranyl diphosphate (GGPP), with the consequent accumulation of unprenylated small GTPase proteins including those of the Rab and Rho superfamilies (Luckman et al., 1998) (Fig 2a). To address whether N-BP has pharmacological effects on AMϕ and PMϕ, we used liquid chromatography tandem mass spectrometry (LC-MS/MS) to examine the accumulation of IPP/DMAPP and a sensitive biochemical *in vitro* assay to detect changes in the level of unprenylated Rab proteins (Ali et al., 2015, Rogers et al., 2020).

**Figure 2.**
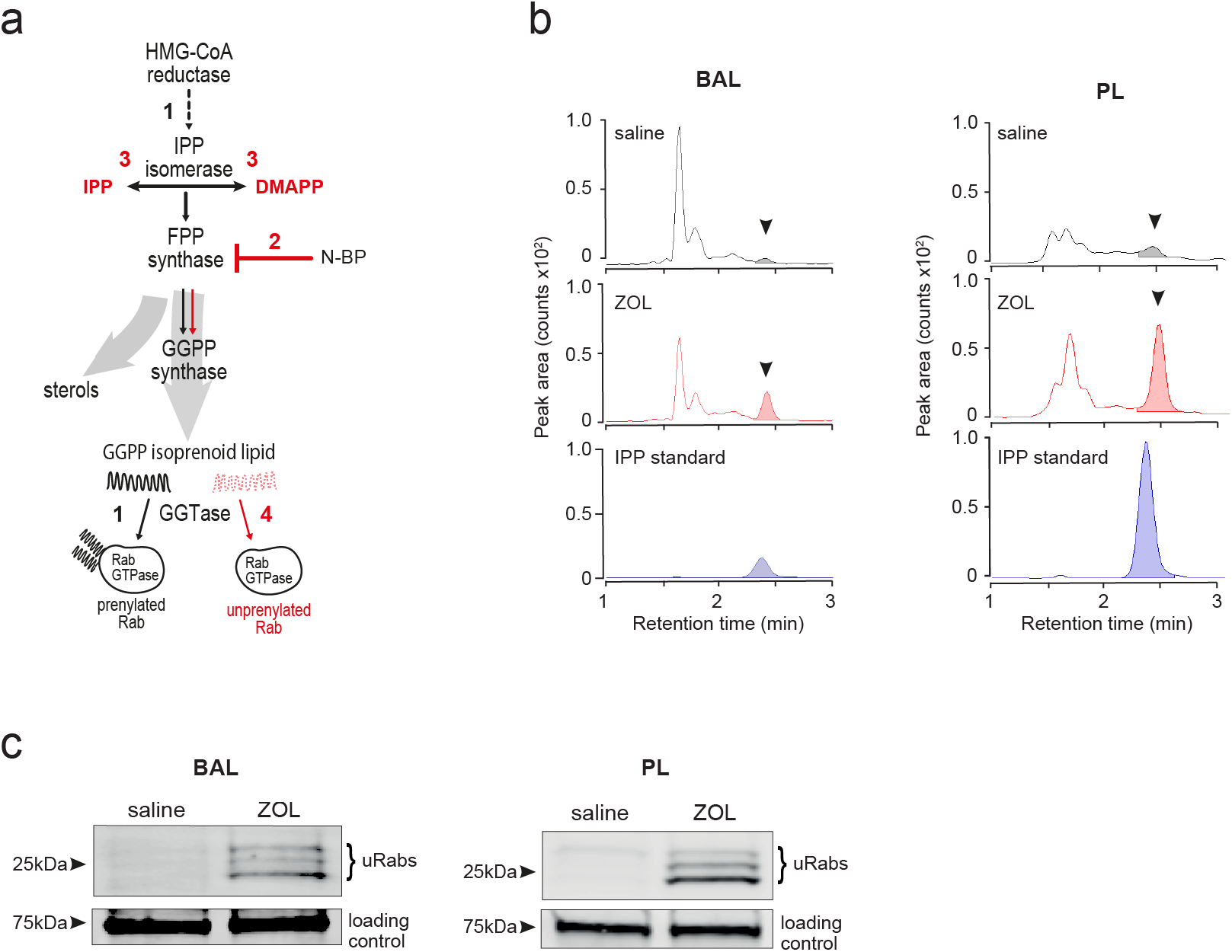
Systemically administered bisphosphonate has pharmacological activity on the mevalonate pathway in tissue-resident macrophages *in vivo.* **a)** Flux though the mevalonate pathway (black arrows) enables prenylation of Rab GTPases by utilising GGPP (step 1); inhibition of FPP synthase by N-BP (step 2) causes upstream build-up of IPP and DMAPP (step 3) and prevents downstream Rab prenylation by reducing GGPP synthesis (red arrows, step 4). **b)** Detection of IPP/DMAPP (arrowhead) by LC-MS/MS in cells from BAL (left) and PL (right) samples 48 hours after *i.v.* ZOL or saline treatment. Coloured peaks in the chromatogram depict the relative abundance of IPP/DMAPP. **c)** Detection of unprenylated Rab GTPases (uRabs) in cells from BAL (left) and PL (right) samples 48 hours after *i.v.* ZOL or saline treatment. Data are representative of 3 separate experiments.

LC-MS/MS analysis showed that IPP/DMAPP was undetectable in extracts of BAL or PL cells from saline-treated mice, but there was a clear increase in the level of IPP/DMAPP in BAL and PL cells collected 48 hours after a single *i.v.* injection of the N-BP ZOL (Fig 2b). Importantly, ZOL treatment also resulted in a marked accumulation of unprenylated Rab proteins in BAL and PL cell samples (Fig 2c). It is unlikely that such an effect of N-BP on protein prenylation in macrophages *in vivo* could have been detected using the relatively insensitive western blot approach previously employed to study bone-resorbing osteoclasts, which engulf large amounts of N-BP (Frith et al., 2001, Rogers et al., 2020). However, the development of a much more sensitive *in vitro* prenylation assay (Ali et al., 2015) now allows the detection of subtle effects on protein prenylation in cells outside the skeleton that may internalise much smaller quantities of N-BP.

Together with the evidence for uptake of N-BP by large PMϕ and AMϕ (Fig 1), these findings demonstrate unequivocally for the first time that systemic administration of N-BP has pharmacological activity outside the skeleton. We show that a single dose of ZOL is sufficient to inhibit the mevalonate pathway in tissue-resident macrophages (AMϕ and PMϕ), causing a build-up of IPP/DMAPP metabolites and an accumulation of unprenylated small GTPase proteins – characteristic hallmarks of N-BP action (Rogers et al., 2020). Although still to be confirmed in humans, our findings overturn the longstanding textbook paradigm that N-BP drugs, which have been in clinical use for several decades (Russell, 2011), act only in the skeleton.

### Treatment with N-BP *in vivo* enhances the production of cytokines and chemokines in response to immune challenge

We recently reported that loss of protein prenylation in cultured monocytes promotes the formation of the NLRP3 inflammasome, resulting in increased caspase-1-mediated processing of pro-IL-1β following bacterial endotoxin (lipopolysaccharide/LPS) stimulation (Skinner et al., 2019). Therefore, we next examined whether N-BP-mediated inhibition of protein prenylation in tissue-resident macrophages alters the response to LPS *in vivo*, particularly in the lung. Mice were challenged intranasally (*i.n.*) or intraperitoneally (*i.p.*) with LPS 48 hours after *i.v.* administration of ZOL or saline (Fig 3a). ZOL treatment alone did not alter cell viability, the ratio or total number of macrophages recovered in BAL samples (Suppl Fig 1b,c), nor had any effect on the levels of cytokines/chemokines in BAL fluid (Fig 3b). However, *i.n.* LPS administration in ZOL-treated mice resulted in a significant increase (2.5-5.0 fold) in the production of proinflammatory cytokines IL-1β, IL-6, TNFα, G-CSF, GM-CSF and chemokines CXCL1, CCL2, CCL3, CCL4 and CCL5 in BAL fluid compared to control mice (Fig 3b). This increase in cytokine and chemokine release was not associated with changes in cell viability, total or relative numbers of BAL cells or AMϕ (Suppl Fig 1b,c).

**Figure 3.**
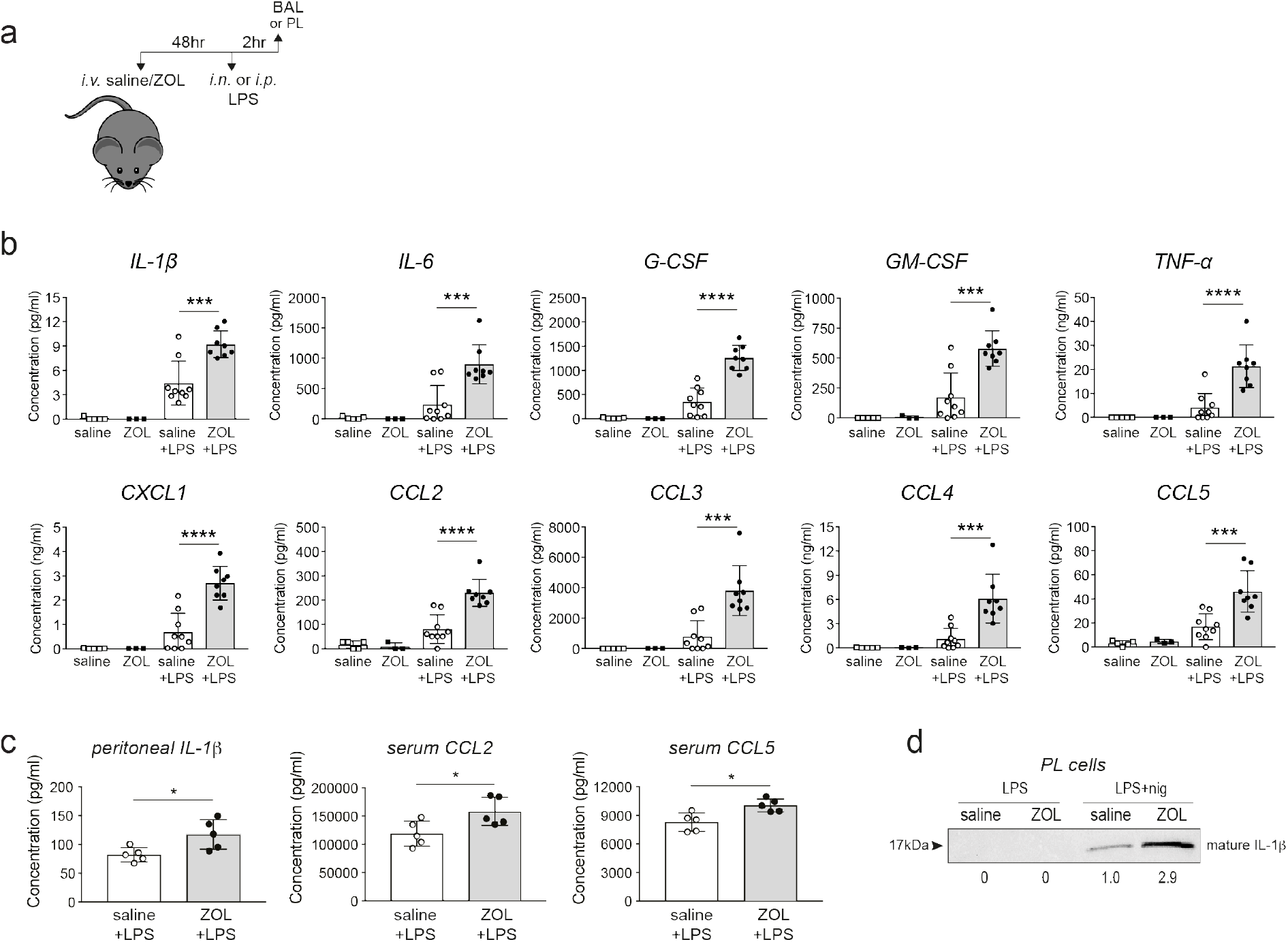
ZOL treatment enhances the production of inflammatory cytokines in response to endotoxin challenge *in vivo.* **a)** Schedule of *i.v.* ZOL administration 48 hours prior to *i.n.* or *i.p.* LPS treatment and subsequent collection of BAL or PL fluid, respectively. **b)** Multiplex analysis of cytokines and chemokines in BAL fluid from saline- or ZOL-treated mice after *i.n* LPS challenge, and **c)** in peritoneal fluid,and serum after *i.p.* LPS challenge. In **b** bars represent mean ± SD, n=8-9 mice per group with LPS, or n=3-5 mice per group with ZOL/saline alone; ***p<0.001, ****p<0.0001, ANOVA with Tukey’s post-hoc test. In **c** bars represent mean ± SD, n=5 mice per group; *p<0.05, unpaired t-test with Welch’s correction; each symbol represents the measurement from an individual mouse. **d**) Western blot detection of mature, extracellular IL-β in conditioned medium from PL cells, isolated from a ZOL- or saline-treated mouse then stimulated *ex vivo* with LPS or LPS+nigericin. Relative levels of IL-β were calculated by densitometry and shown below each lane. The blot shown is representative of three independent experiments.

*I.p.* LPS challenge of ZOL-treated mice also resulted in a significant elevation in IL-1β in peritoneal fluid, and CCL2 and CCL5 in serum, compared to controls (Fig 3c). Furthermore, peritoneal cells from ZOL-treated mice produced almost 3-times more IL-1β upon *ex vivo* stimulation with LPS and nigericin (a well-described NLRP3 activator) (Fig 3d), than cells from control mice. IL-1β is primarily produced by monocyte/macrophages (Kany et al., 2019), the predominant cell type in PL (Suppl Fig 1a). Thus, our results strongly suggest that macrophages are the most likely source of IL-1β. Importantly, treating the mice with ZOL did not cause IL-1β release from peritoneal cells stimulated *ex vivo* with LPS in the absence of nigericin (Fig 3d), and this is in accord with our previous finding that inhibition of the mevalonate pathway alone does not trigger NLRP3 inflammasome assembly but enhances its activation (Skinner et al., 2019).

The observations described here begin to provide a plausible mechanistic explanation for the decreased risk of pneumonia mortality associated with N-BP treatment (Colon-Emeric et al., 2010, Sing et al., 2020, Reid et al., 2021). AMϕ are the initial line of defence against common respiratory tract infections (Byrne et al., 2015), and inhibition of the mevalonate pathway in these cells may help boost the initial response to bacterial as well as viral lung infections by a variety of routes (Fig 4). *First*, uptake of N-BP into cells causes the accumulation of IPP/DMAPP (Fig 2b), which activates human Vγ9Vδ2-T cells (Thompson and Rogers, 2004). Vγ9Vδ2 are non-conventional T cells with potent anti-bacterial and anti-viral activity that recognise phosphoantigens, including IPP, derived from the mevalonate or DOXP pathways in bacterial pathogens (Tanaka et al., 1995, Jomaa et al., 1999). There is considerable interest in the use of N-BP-expanded γ,δ-T cells as an immunotherapy for cancer (Clezardin and Massaia, 2010, Tanaka, 2020) as well as viral diseases (Juno and Kent, 2020). Indeed, N-BP-expanded Vγ9Vδ2-T cells reduce disease severity and mortality from influenza A virus (IAV) infection in humanised mice (Tu et al., 2011, Zheng et al., 2015). *Second*, the genome of some pathogens such as IAV encodes proteins with a prenylation motif, which require the host cell’s mevalonate pathway to enable prenylation and allow pathogen propagation (Marakasova et al., 2017). Agents that block the mevalonate pathway (such as statins) or that inhibit prenylation (such as lonafarnib, currently in clinical trials for hepatitis delta virus infection), have well-described anti-viral or anti-microbial effects (Parihar et al., 2019, Einav and Glenn, 2003). Intriguingly, simvastatin was shown to improve outcomes in hospitalised older adults with community-acquired pneumonia (Sapey et al., 2019). *Third*, inhibition of FPP synthase by N-BP in AMϕ mimics the decreased flux through the mevalonate pathway in macrophages in response to endogenous IFN signalling, which serves to limit viral uptake and replication by several mechanisms including the synthesis of 25-hydroxycholesterol (Robertson and Ghazal, 2016, Cyster et al., 2014). *Fourth*, lack of protein prenylation (Fig 2c) enhances NLRP3 inflammasome activation and promotes the release of IL1-β (Skinner et al., 2019). IL1-β is a central mediator of the innate immune response that orchestrates the production of a cascade of cytokines and chemokines (Garlanda et al., 2013). Our observation that systemic ZOL treatment significantly enhanced the release of IL-1β and several other cytokines and chemokines in lung, peritoneum and serum after LPS challenge (Fig 3), is consistent with increased inflammasome activation. To our knowledge there is no evidence that enhanced cytokine production caused by N-BP therapy worsens lung inflammation in pneumonia patients – on the contrary, N-BP treatment appears to have a beneficial effect on pneumonia risk and mortality (Colon-Emeric et al., 2010, Sing et al., 2020, Reid et al., 2021).

**Figure 4.**
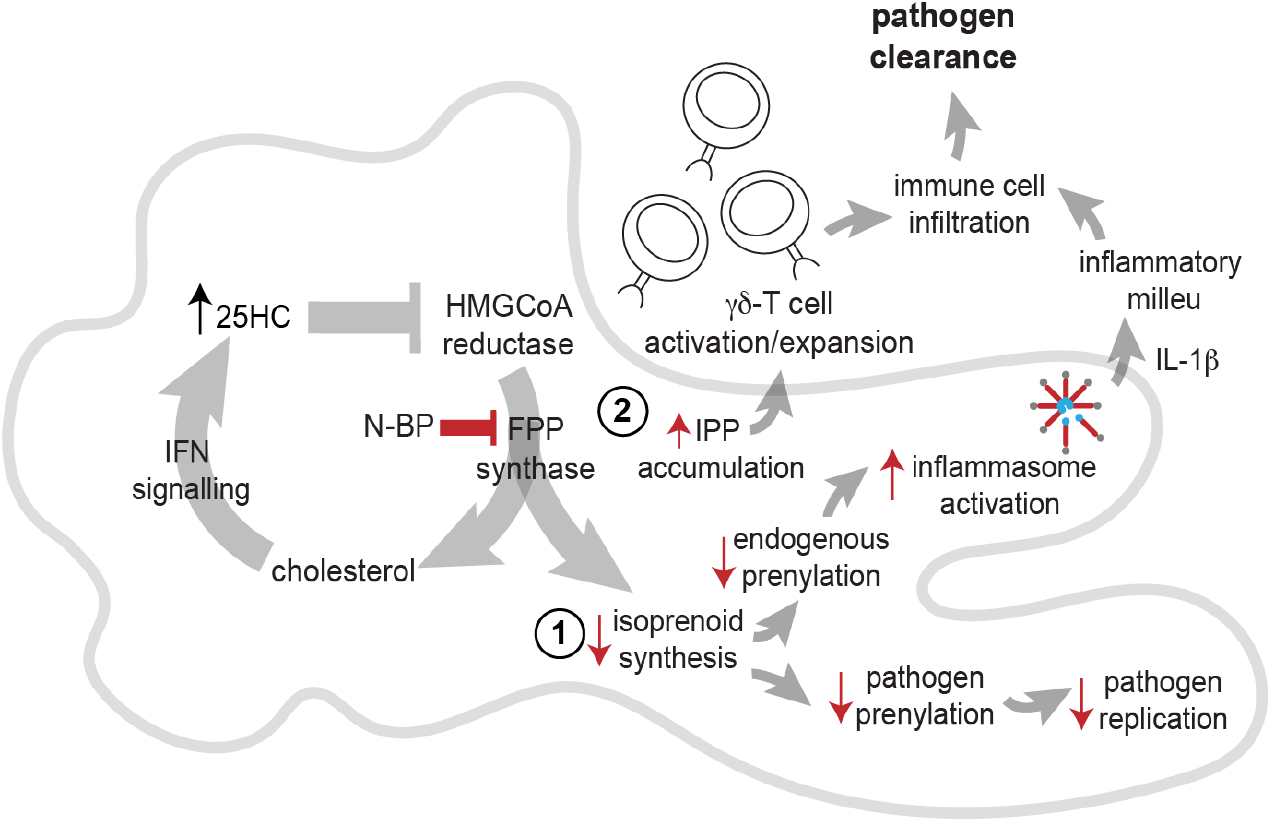
Potential routes of antimicrobial activity of N-BP *via* effects on the mevalonate pathway in alveolar macrophages. Inhibition of FPP synthase by N-BP prevents the biosynthesis of isoprenoid lipids required for normal protein prenylation (step 1). Lack of prenylation leads to enhanced inflammasome activation and increased IL-1β release in response to bacterial endotoxin, boosting the initial inflammatory response and pathogen clearance. Lack of isoprenoid biosynthesis may also hinder the propagation of intracellular pathogens that depend on the host cell’s mevalonate pathway. Inhibition of FPP synthase by N-BP causes the accumulation of IPP/DMAPP phosphoantigens (step 2) capable of triggering activation and proliferation of human Vγ9Vδ T cells with anti-microbial activity. Inhibition of FPP synthase by N-BP may also mimic the endogenous anti-viral effect of 25-hydroxycholesterol (25HC), one of the routes by which IFN signalling suppresses the flux through the mevalonate pathway.

Finally, it is noteworthy that ZOL was recently identified, using computational biology approaches, as one of 200 clinically-approved drugs that are predicted to target pathways induced by SARS-CoV-2 and could be suitable for drug repurposing against COVID-19 (Han et al., 2021). Epidemiological studies are therefore urgently needed to determine whether N-BP therapy alters the incidence, severity or risk of mortality from SARS-Cov-2 infection.

## CONCLUSIONS

We show that systemically administered N-BP drug can inhibit the mevalonate pathway and prevent protein prenylation in tissue-resident macrophages beyond the skeleton, and this in turn enhances macrophage responsiveness to bacterial endotoxin. Our observations in mice, together with data from clinical studies in humans (Colon-Emeric et al., 2010, Sing et al., 2020, Reid et al., 2021), suggest that the beneficial effects of these drugs against pneumonia infection and mortality are, at least in part, mediated by targeting AMϕ, thereby boosting early immune responses in the lung. These findings are particularly relevant to the elderly population, and clinical trials are warranted to determine whether N-BP drugs, aside from preventing bone loss, can provide protection to pneumonia infection in vulnerable individuals, for instance, patients in aged care homes.

## MATERIALS AND METHODS

### Animals and tissue collection

Studies involving mice were performed in strict accordance with the Australian Code for the care and use of animals for scientific purposes (2013). All of the animals were handled according to Animal Ethics Committee protocols (Animal Research Authority: 18/40) approved by the Garvan Institute/ St Vincent’s Hospital Animal Ethics Committee. Procedures were performed under appropriate anaesthesia, with animal welfare consideration underpinned by the principles of Replacement, Reduction and Refinement.

All experiments involved female C57BL/6J mice and group sizes were based on previous studies using the same methodologies. Animals were purchased from Australian BioResources, housed with standard chow diet in specific pathogen-free conditions and randomly allocated to experimental groups. Mice were anaesthetised with isoflurane prior to retro-orbital intravenous (*i.v*.), intranasal (*i.n*.) or intraperitoneal (*i.p.*) drug administration. Mice were euthanised by CO_2_ inhalation and peritoneal and bronchoalveolar cells were isolated by lavage using 2mM EDTA/magnesium- and calcium-free DPBS (Gibco). Peritoneal lavage (PL) cells were collected by injecting 5mL of solution into the peritoneal cavity, and bronchoalveolar lavage (BAL) cells by pooling 3 consecutive 1.5 mL washes administered *via* an insertion in the trachea.

### Flow cytometry

A single dose (47.5 μg /40 nmoles) of AF647-ZOL (BioVinc, CA, USA), or 500μg/kg ZOL (or saline vehicle) were administered *i.v.* in a final volume of 100 μL. Uptake of AF647-ZOL was assessed in BAL and PL cells 4 hours later. For LPS challenge, mice were treated with *i.v*. ZOL 48 hours prior to administration of *i.n*. LPS (10 μg LPS in 20μL saline) or *i.p*. LPS (100 μg in 200μL saline). Cell viability and total cell numbers in BAL and PL were assessed by trypan-blue staining using a Corning CytoSMART cell counter. For flow cytometric analysis, cells were pre-incubated with mouse-Fc block and viability marker prepared in calcium/magnesium-free PBS (Gibco), before staining with fluorescently-conjugated antibodies prepared in staining buffer (2mM EDTA, 0.02% azide; 0.5% foetal calf serum, in calcium/magnesium-free PBS). Samples were analysed using a BD LSRII SORP flow cytometer/ DIVA software. The post-acquisition analysis was performed using FlowJo 10.6.2 (BD).

### Antibodies

Anti-CD11b-BUV395 clone M1/70 (1:200 dilution); anti-MHCII-FITC clone 2G9 (1:200); anti-CD11c-PE clone N418 (1:200); anti-CD11c-BV421 clone HL3 (1:200); anti-B220-BUV737 clone RA3-6B2 (1:200); anti-Ly6C-PE-Cy7 clone AL-21 (1:200); anti-TCRB-APC-Cy7 clone H57-597 (1:300); anti-F4/80-BV650 clone BM8 (1:100); anti-Siglec-F-BB515 clone E50-2440 (1:200); mouse-Fc block clone 2.4G2 (1:200); and streptavidin-BV421 (1:400) were purchased from BD Biosciences. Anti-F4/80-Biotin clone BM8 (1:200); anti-Ly6G-APC clone IA8 (1:200); anti-Ly6G-PerCP-Cy5.5 clone IA8 (1:200); and Zombie Aqua viability stain (1:700) were from Biolegend.

### LC-MS/MS analysis of IPP/DMAPP

10-week-old female mice were administered a single 100 μL retro-orbital *i.v.* dose of 500 μg/kg ZOL (or saline control). Animals were culled 48 hours later and broncho-alveolar cells and peritoneal cells were collected by pooling the BAL or PL lavages from n=8 mice (BAL) or n=5 mice (PL). Cell pellets were then stored at −80°C. For analysis of IPP/DMAPP, 1mL cold extraction solvent (80:20 methanol:water) was added to the cell pellets then vortexed for 10 seconds and incubated in a ultrasonic bath filled with ice water for 1 hour, then centrifuged at 3000 rpm for 30 minutes at 4°C. Aliquots (850μL) of supernatant were dried under vacuum in an Eppendorf Concentrator Plus then reconstituted in 42.5μL 70% methanol, 30% 10mM ammonium acetate. Samples (injection volume 1 μL) were analysed by targeted LC-MS/MS using an Agilent 1290 Infinity II UHPLC system coupled to an Agilent 6495B triple quadrupole mass spectrometer. Separation was achieved using an Agilent Infinity Poroshell 120 EC-C18 column (3.0×150mm, 2.7μm) fitted with an Agilent Infinity Poroshell 120 EC-C18 UHPLC guard column (3.0×150 mm, 2.7 μm), maintained at 20°C. The mobile phases were 10 mM ammonium acetate in water (A) and methanol (B), both containing 5 μM medronic acid to chelate metal ions (gradient 98% A from 0 to 3 minutes, decreased to 2% A from 3.5 to 6.5 minutes at 0.5 mL/min, then increased to 98% A at 0.4 mL/min from 6.5 to 12 minutes (total run time 12 minutes). Autosampler temperature was 4°C. The mass spectrometer was operated in negative electrospray ionisation mode: source gas temperature was 250°C with flow at 17 L/min, sheath gas temperature was 400°C with flow at 12 L/min, and nebuliser pressure was 45 psi. Data were acquired in MRM (Multiple Reaction Monitoring) mode and was processed using Agilent MassHunter Quantitative Analysis software version B08.00.00. By comparison with pure standard compounds (Sigma Aldrich), the isomers IPP and DMAPP eluted at the same retention time (approximately 2.4 minutes) and were calculated as total area under the curve. The limit of detection in cell extracts was 20nM.

### Detection of unprenylated Rab proteins

To assess the accumulation of unprenylated Rab GTPase proteins we used an *in vitro* prenylation assay as previously described (Ali et al., 2015). Mice were treated with *i.v.* ZOL or saline as described above, then cell pellets were obtained by BAL or PL (each pooled from n=5 mice) and lysed by sonication in prenylation buffer (50 mm HEPES, pH 7.2, 50 mm NaCl, 2 mm MgCl2, 100 μm GDP, 1× Roche complete EDTA-free protease inhibitor cocktail). For the *in vitro* prenylation assay, 10 μg of protein were incubated with recombinant GGTase II, REP-1 and biotin-conjugated GPP (a synthetic isoprenoid lipid) for the labelling of unprenylated Rab proteins(Ali et al., 2015). The resulting biotinylated Rabs were then detected on PVDF blots using streptavidin-680RD (LiCOR). A narrow doublet (often appearing as a broad singlet) of endogenous biotinylated 75 kDa proteins were used as a sample loading control.

### Immune responses to LPS in vivo

10-week-old female mice were administered a single retro-orbital *i.v.* dose of 500 μg/kg ZOL (or saline control), 48 hours before immune challenge with LPS (*E.coli* O111:B4, Sigma-Aldrich) administered either *i.n*. (10 μg LPS in 20μL saline) or *i.p*. (100 μg in 200μL saline). Mice were culled 2 hours and BAL or PL fluid were collected by injecting 500 μL or 1mL PBS into the lungs or peritoneal cavity, respectively. Cytokines and chemokines in BAL, PL and serum were measured using a Bio-Plex multiplex immunoassay (Bio-Rad) and a MAGPIX (Luminex) multiplex reader according to the manufacturer’s instructions.

### IL-1β release by peritoneal cells ex vivo

PL samples were obtained from ZOL- or saline-treated mice as described above. PL cells were placed in 96-well plates (400,000 cells/well) and treated at 37°C for 5.25 hours with 200 ng/ml LPS, in a final volume of 200 μL serum-free Opti-MEM medium (Gibco), followed by stimulation for 45 minutes with 10 μM nigericin (Sigma-Aldrich). The level of mature (17 kDa) IL-1β in conditioned medium was analysed by western blotting on nitrocellulose membrane with a goat anti-mouse IL-1β antibody (AF-401-NA, R&D systems, 1:1000 dilution), donkey anti-goat horseradish peroxidase-conjugated secondary antibody (A15999, Thermo Fisher, 1:5000 dilution) and enhanced chemiluminescence using SuperSignal West Pico chemiluminescent substrate (34580, Thermo Fisher). The signal was detected using a Fusion FX7 imaging system (Etablissements Vilber Lourmat SAS) and densitometry was performed on blots using ImageJ (v2.0.0).

## ACKNOWLEDGEMENTS

We thank Prof Kirill Alexandrov (Queensland University of Technology) and Dr Zakir Tnimov (MRC Laboratory of Molecular Biology) for providing reagents for the Rab prenylation assay. This work was supported in part by National Health and Medical Research Council (NHMRC) of Australia project grant 1079522 to MJR, by Mrs Janice Gibson and the Ernest Heine Family Foundation, and a Perpetual IMPACT grant to MAM. PMH is funded by a Fellowship and grants from the NHMRC (1175134) and by University of Technology Sydney (UTS). We gratefully acknowledge funding by the New South Wales Government for the Victor Chang Cardiac Research Institute Innovation Centre, as well as funding from the Freedman Foundation for the Metabolomics Facility.

## COMPETING INTERESTS

FHE is an employee of BioVinc (Pasadena, CA, USA). SS also co-affiliates with BioVinc. JRC participates on advisory boards of Amgen and Bayer and receives payment from Amgen for educational activities. The other authors declare no competing interests.

---

**Supplementary Figure 1.**
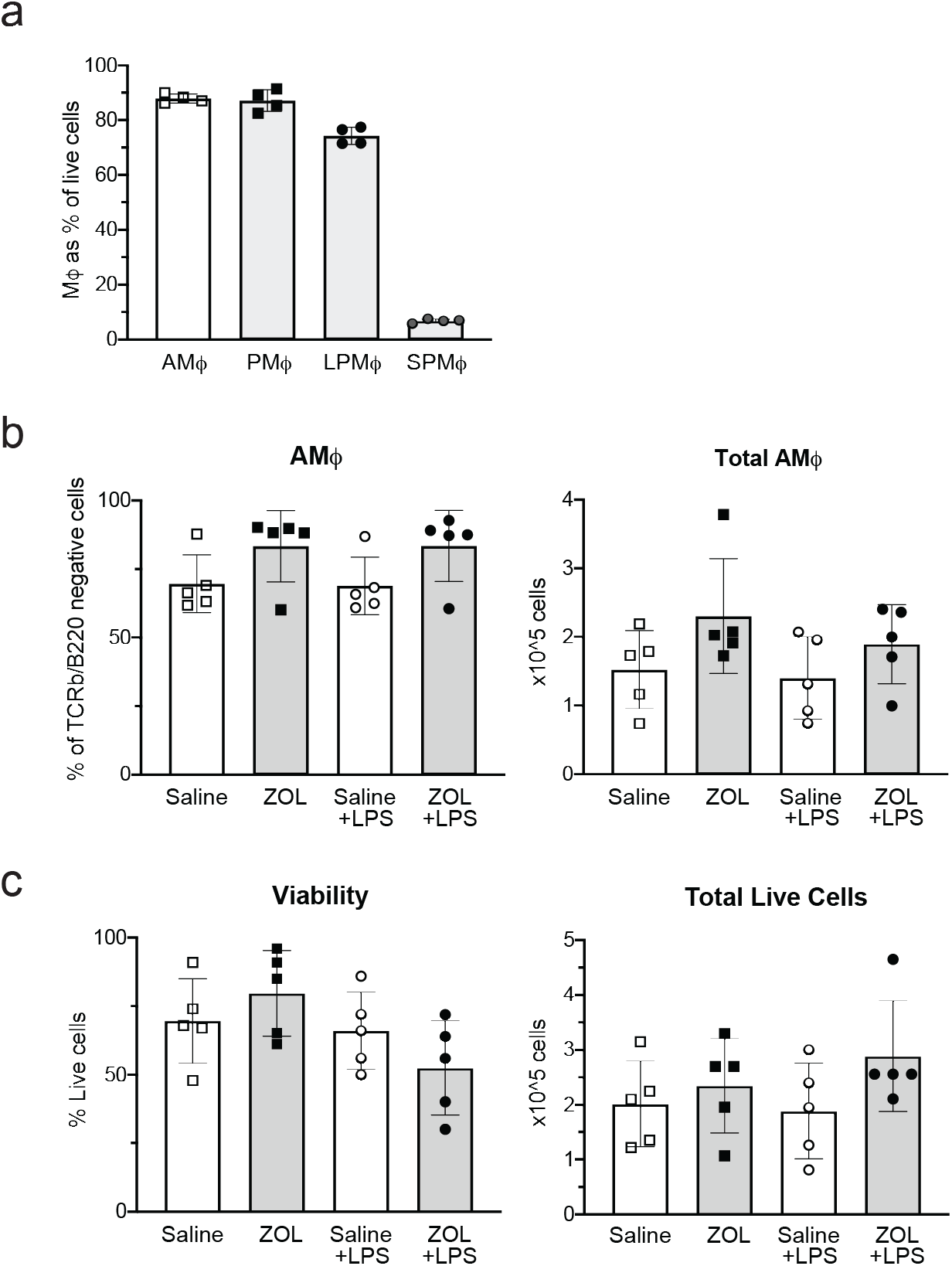
ZOL treatment does not affect the number or viability of tissue-resident macrophages *in vivo*. Flow cytometric analysis of cells in BAL and PL samples from saline-treated mice (n=4 mice per group). The graph shows the percentage of AMϕ (alveolar macrophages: B220^−^TCRb^−^ CD11c^hi^F4/80^+^Siglec-F^hi^ singlets) and PMϕ (total peritoneal monocyte/macrophages: B220^−^TCRb^−^Siglec-F^−^, Ly6G^−^ singlets), LPMϕ (large peritoneal macrophages: B220^−^TCRb^−^Siglec-F^−^ Ly6G^−^CD11b^hi^F4/80^hi^ singlets) and SPMϕ (small peritoneal macrophages: B220^−^TCRb^−^Siglec-F^−^Ly6G^−^CD11b^lo^F4/80^lo^ singlets). **b)** Percentage (left) and total cell number (right) of AMϕ in BAL as analysed by flow cytometry. **c)** Viability (left) and total cell number (right) in BAL measured by trypan blue exclusion. Samples in **b** and **c** were obtained from mice injected *i.v* with saline (white bars) or ZOL (grey bars) 48 hours prior to *i.n.* administration of saline (square symbols) or LPS (round symbols) (n=5 mice per group). Bars represent mean ± standard deviation; each symbol represents the measurement from an individual mouse.

## LEGEND TO SOURCE DATA

Files contain TIF images of protein blots, after *in vitro* prenylation of BAL or PL cell lysate samples to detect unprenylated Rab GTPases (uRabs), or western blot of conditioned media to detect IL-1β:

**Figure 2c BAL_source data 1,2**

**Figure 2c PL_source data 1,2**

Following the *in vitro* prenylation assay (described by Ali *et al* 2015, *Small GTPases* **6**, 202-211) blots typically show a cluster of three bands around 23-27 kDa (unprenylated Rab proteins) and several bands of endogenous biotinylated proteins including around 40 kDa, 120 kDa, and a band around 75 kDa of an endogenous, ubiquitously expressed protein used as a loading control (Ali *et al*, *Small GTPases* 2015). Brightness and contrast of the entire blot was adjusted in Image Studio software (LiCOR) before cropping areas of uRabs and the loading control band.

**Figure 3d_source data 1,2**

Western blot of conditioned media using anti-IL-1β shows a single 17 kDa band of cleaved IL-1β.

